# Semi-analytical dynamic modeling of DNA surface-hybridization via AC Electro-kinetic steering

**DOI:** 10.1101/2020.02.25.964809

**Authors:** P. Capaldo, S. D. Zilio, V. Torre, Y. Yang

## Abstract

The change in electrical property (capacitance) upon hybridization of the desired ssDNA to a capture probe has been proposed as a promising technology platform in biomedical research and practice. An appropriate mathematical model is needed for understanding and optimizing the process occurring at the electrode/electrolyte interface. It is also informative for examining the forces generated by the AC electric fields on the DNA molecules as well as the suspending buffer solution in the experimental pool. Here, we provide the development, formulation and validation of a semi-analytical model of DNA hybridization with deoxynucleotide molecules chemically tethered to a solid gold electrode. The parameters of the proposed model have been estimated using available experimental data. We demonstrate that the detection limit and specificity of our surface-based genosensor are not only dependent on the probe/target binding affinity, but also on the Self-Assembled Monolayer (SAM) density and on the interfacial electric field. The label-free Electrochemical Impedance Spectroscopy (EIS)-based oligonucleotide biosensor with integrated DC-biased can achieve rapid hybridization, high selectivity and sensitive detection for DNA target samples.

**SIGNIFICANCE:** DNA hybridization, wherein strands of DNA form duplex through noncovalent, sequence-specific interactions, is one of the most fundamental processes in biology. Fast and reliable determination of miniature amounts of DNA plays important role in clinical forensic and pharmaceutical applications. Thus, developing a better understanding of the kinetic and dynamic properties of DNA hybridization will help in the elucidation of all mechanisms involved in numerous biochemical processes. Moreover, because DNA hybridization has been widely adapted in biotechnology, its study is invaluable to the development of a range of commercially important processes.

To achieve optimal sensitivity with minimum sample size and rapid hybridization, ability to predict the kinetics of hybridization based on the characteristics of the strands is crucial, and hence a computer aided numerical model for the design and optimization of a DNA biosensor has been implemented.

## INTRODUCTION

DNA is central to biology and has become a key ingredient in nanotechnology. The determination of specific nucleic acid sequences originating from organisms has attracted widespread interest for use in medical diagnostics such as the identification of pathogenic species and the recognition of genetic diseases, drug discovery, food safety monitoring and agriculture.

In particular, label-free methods promise easy, fast and cost-effective DNA detection. Among the label-free methods, EIS-based biosensors exhibit several advantages, including inherent high sensitivity, good compatibility with electrical instruments and ease of large-scale production, making them well-suited for diagnostic systems (1–3).

Although label-free DNA biosensors can significantly simplify oligonucleotide detecting procedures, the hybridization process still needs long time to reach a plateau in stationary solutions. The limiting step is mainly due to the Brownian motion of nanometer-scale targets, moving stochastically toward the probes immobilized on the sensor surface. Furthermore, the diffusion is influenced by a number of factors, and in particular by the electrode surface properties and by the number of active and available probe sites. Therefore, electrode design requires in-depth knowledge of the micro-flow mass transport phenomena and their effects on the performances of the sensor.

It has been shown that the effectiveness of DNA sensors and biosensors depends on the accuracy of the prediction of the experimental parameters responsible for the thermostability and for the time of the DNA duplexes forming. As for all-natural processes, solid-phase hybridization is characterized by thermodynamic and kinetic restrictions: the thermodynamics acts as discriminating factor for the different target sequences and detection of low concentration targets, while the kinetics for how quickly the equilibrium is approached.

The thermodynamics of DNA duplex formation is dominated by reactions involving either bound duplexes or separated strands, which has been deeply investigated, experimentally and theoretically (all-or-nothing two state models) (4–6). By contrast, DNA hybridization kinetics depends on the intermediate states occurring between two steady states, which are transient and complicated processes, and are therefore difficult to investigate and measure quantitatively

Several groups have examined DNA hybridization by theoretical approach using a variety of strategies. The thermodynamics and kinetics of hybridization have been thoroughly studied, both in the bulk and at the surface. Chan et al. (7) and Erickson et al. (8) considered the case in which DNA probes are immobilized at low density, permitting surface adsorption and lateral diffusion of DNA targets. Hagan and Chakraborty (9) explored the effect of steric crowding on initial hybridization rate constants using polymer brush models. Wong and Melosh (10) modeled experimental results at high DNA densities taking into account the effects of applied potential on the biomolecular activation and the changing electrostatics within the layer, which rapidly becomes nonlinear as hybridization proceeds (11,12).

Although numerous experimental studies of the kinetics of oligonucleotide hybridization have been reported, a theoretical analysis of the influence of an alternating potential on the DNA hybridization mechanism is necessary.

## METHODS

The present work is devoted to develop a semi-analytical model for the estimation of the rate constants of non-competitive oligonucleotides hybridization in a pure salt solution (100 mM KCl) and on a gold-modified surface, in connection to a capacitive genosensor development.

Based on our previous works (13–15), we developed a miniaturized, label-free, electrochemical impedance spectroscopy-based device with which we were able to perform *real-time*, and *in-situ* detection of double layer capacitance, C_DL_, changes occurring at the functionalized interface, see Supporting Materials, Section II, for more details.

C_DL_ variations, depend mainly by the height increase of Self-Assembled Monolayer (SAM), adsorbed on the electrode surface, due to the different persistence length of ssDNA and dsDNA; and by the replacement of water molecules with DNA molecules on DNA pairing resulting in changes in the electrical charge density.

In our operating conditions, C_DL_ is equal to the differential capacitance, C_d_, at the electrode/electrolyte interface and can be modeled, in first approximation, as a conventional capacitor. We can compute the corresponding differential capacitance as:

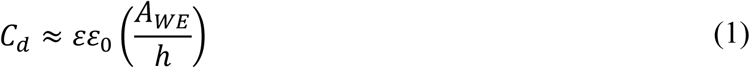

where A_WE_ (cm^2^) is the working electrode area, h is the thickness, expressed in nm, of molecular layer, ε_0_ and ε are the vacuum permittivity and molecular layer dielectric constant, respectively.

Therefore, measurement of the SAMs’ thickness and compactness provide insights into our system modelling. Tarlov and co-workers (16, 17) pioneered a technique for DNA immobilization that utilizes the chemisorptive self-assembly of thiol-terminated single-stranded DNA (HS-ssDNA) monolayers onto gold surfaces, mixed with a short hydroxyl-terminated alkylthiol surface diluent [e.g., mercaptohexanol (MCH)] to prevent the nonspecific attachment of DNA to the surface by nucleotide amine groups and enhance specific attachment by the thiolated group.

In this work, quantitative atomic compositions of the mixed DNA monolayers were estimated using X-ray Photoelectron Spectroscopy (XPS) and a more detailed description is provided in Supporting Material, Section III.

Given a SAM density of *ρ*_*SAM*_ ≃ 2 × 10^12^ molecules cm^−2^ (13) as stated by Peterson et al. (18), the DNA hybridization efficiency can reach about 100%. However, in this *pseudo-Langmuir (PL) regime*, the steric crowding effect on initial hybridization rate was not considered. Besides, at our ionic composition and concentration, and SAM-density, we cannot neglect probe-probe interactions (as provided in Supporting Material, Section VIII, Eq. S16) that affect the hybridization reactions leading to a lower number of available sites and, hence, to a kinetic deletion of hybridization.

In this framework, Wong and Melosh (10) quantified hybridization target number density as a function of probe coverage and ion concentration; they demonstrated that, in a *suppressed hybridization regime*, the electrostatic potential, outside the DNA probe layer, reduces the hybridization efficiency by repelling incoming DNA targets. (see the paper for more details).

In the experimental conditions (i.e. ionic strength, [cDNA] in solution) similar to that of our study, they found an efficiency of hybridization of about 50%. In addition, the characteristic timescale needed to hybridize 1 µM of [cDNA] is ∽ 400 s, consistent with Georgiadis group’s measurements (18). This hybridization time is mainly due to the stochastic Brownian motion by which cDNA strands move towards the surface-immobilized probe ssDNA. Although this timescale is not experimentally prohibitive, lower values [cDNA] may have significantly longer hybridization times going to adversely affect the detection.

To improve the hybridization rate, an alternating current (AC) electric field (see Supporting Material for a detailed description of the role played by the applied voltage in the increase of DNA hybridization efficiency) and other electrokinetiks mechanisms such as dielectrophoresis (DEP) and electroosmotic flow (ACEO), can be used to manipulate fluids and biomolecules (smaller than 1 µm in diameter) without the use of external pumps (19–21).

**DEP** arises from a difference in electrical permittivity between the molecule and the surrounding medium. If DEP is positive, the molecule is more polarizable than the medium and the DEP force is directed towards regions of high electrical field strength. By contrast, if the molecule is less polarizable than the buffer solution (DEP < 0) the force is directed down the electric field. Although the DEP force can concentrate several thousand of base double-strand DNA (dsDNA) (22, 23), can not be used to directly manipulate tens of base ssDNA at a nanometer-scale size because proportional to particle volume.

In our working conditions (i.e. single strand complementary 22-mer DNA in 100 mM KCl-doped aqueous solution), the time-averaged DEP is proportional to *Re*{*K*(*ω*)} ≈ −0.223 < 0, which means that when manipulating short and single DNA strands, the DEP force is insufficient to hold the molecules down over the electrode and the electroosmotic drag should be dominant. More details on the time-averaged DEP force estimation can be found in SI.

**AC electroosmosis** (Fig. 1 A) is a flow motion that occurs due to the interaction of the electrolyte ions in the double layer with the transverse component of the electric field (Cartoon in Fig. 1 B). ACEO has the great advance to create a solution vortex in our microsystem and hence improve the hybridization efficiency of tens of base DNA strands. Hart et al. (24), for example, have used the AC-electroosmosis to enhance the transport of analyte to an interdigitated planar electrode, founding a binding time reduced by up to a factor of six.

**FIGURE 1.**
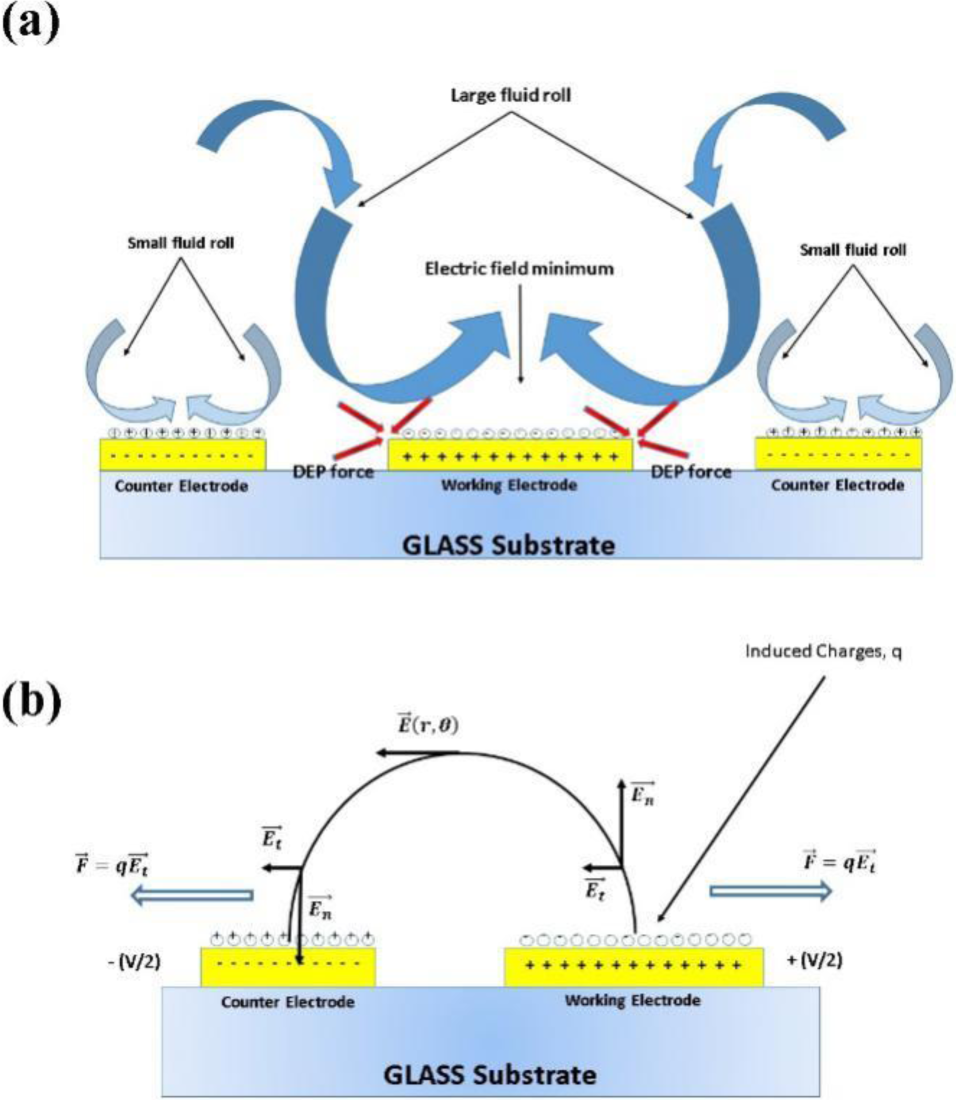
Side-view schematic of rotating vortices induced by AC-ElectroOsmosis above the electrodes. The caused fluid motion is shown as blue arrows as well as red arrows indicate the force due to DEP. Noteworthy, the DNA-SAM film is exposed to a field gradient at the metal/electrolyte interface of approximately 10^9^ V/m.

However, a complete computation of an AC electroosmotic flow would require coupled simulations of the electric field, charge density, conductivity, and fluid velocity within the electrical double layer, as well as computations of the electric field and fluid velocity in the bulk of the device. Henceforth, the simulation strategy proposed by Gonzalez and coworkers (25, 26) was used.

This approach relies on a linear capacitance approximation for the double layer and a fluid slip boundary condition just outside the double layer itself. In addition, it can be applied to electrodes immersed in a symmetric electrolyte with a constant conductivity within the double layer (see SI for more details).

In these assumptions, the time-averaged slip velocity, <u_slip_>, is derived from the Helmoltz - Smoluchowski formula (27–29):

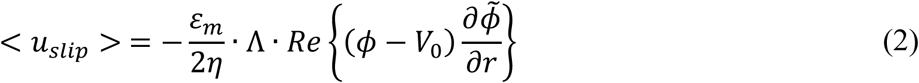

where V_0_ is the applied voltage, ε and η are the permittivity and viscosity of the medium (both assumed constant). Λ is a tuning parameter to account for the Stern layer and it is defined as the ratio of the diffusive-layer to compact-layer capacitance and it is taken equal to 0.25 in the simulation (26), and ϕ is the electrical potential.

The ACEO velocities are implemented and simulated using the computer algebra program Wolfram Mathematica ver. 10 that allow us to compute a FEA of the problem.

The results shown in Fig. 2 indicate that the ACEO applied at the asymmetric CE/WE electrodes, produces a unidirectional flow from the narrower Counter to the wider Working and its velocity decreases with the distance from the electrode surface due to the reduced tangential electric field away from the WE (26, 30).

**FIGURE 2.**
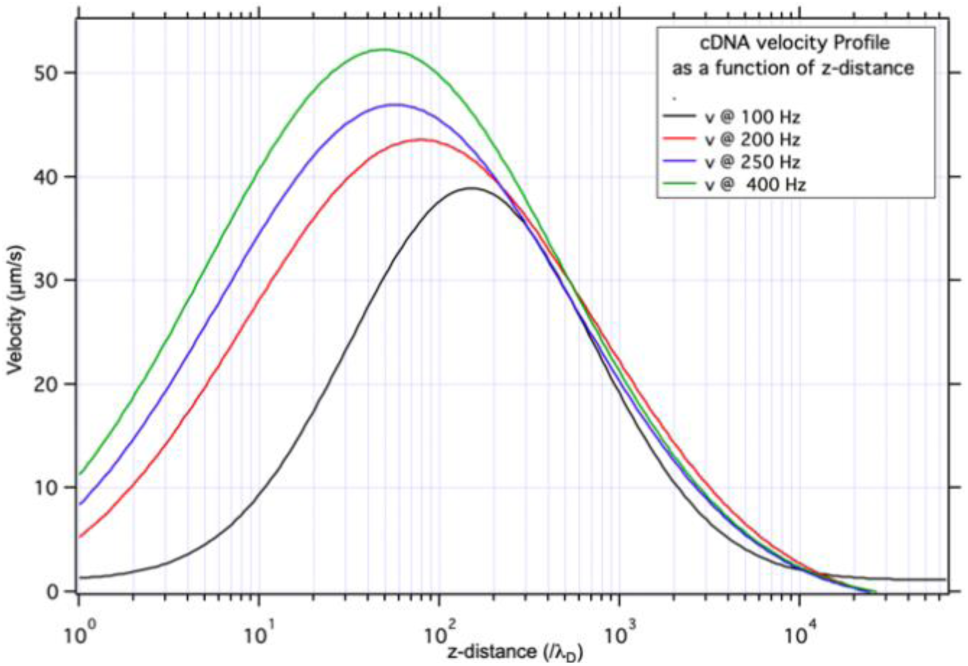
Radial ACEO velocity estimated at 100, 200, 250 and 400 Hz as a function of the distance from the WE surface with V_app_ = 10 mV in a 100 mM-doped aqueous solution and a radius set equal to *R*_*g*_ ≈ 1.78 nm.

The results show that cDNA strands, suspended above the interelectrode gap were attracted to the WE surface (Figs. 3). Thus, following ACEO fluid motion, the cDNA targets are transported towards the center of the working electrode where the electric field has its minimum as a consequence of the negative contribution of DEP forces. The radial velocity (Fig. 3) is computed to be approximately 16 µm/s for an applied voltage of 10 mV peak to peak, a frequency of 250 Hz, electrical conductivity of σ = 55 µS/cm and within a z-distance equals to its dynamic polarization length (31), |*σz*_*H*_| ≈ 1,045 nm, the measure of distance up to which reorganization of charges in solution takes place.

**FIGURE 3.**
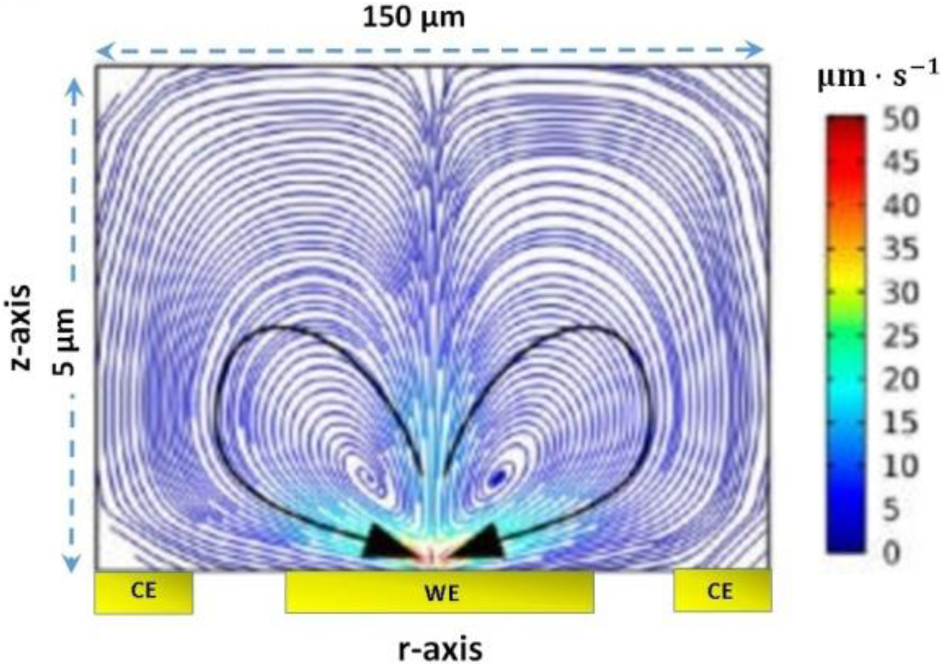
Illustration of the AC electroosmosis fluid flow of cDNA strands in our Immunosensor. The simulation was carried out with an applied voltage of 10 mV at a frequency of 250 Hz. Streamline and arrow plot of electroosmosis flow above working and counter electrodes surface. The colors are plotted on a log scale in units of 10^−6^ m/s.

The velocity profile can be integrated to yield the mass carried by convention into our dynamic polarization length. Into our control volume, the scaling convective flux rate is:

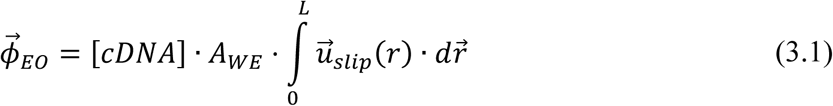

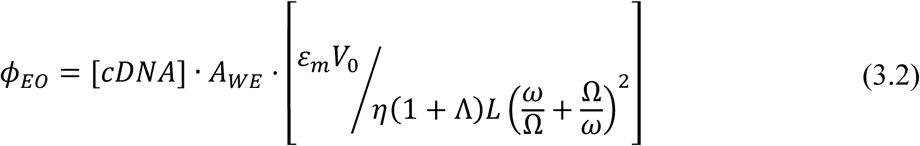

where L is a scale characteristic of the experiments that reflects the changes in the potential along the electrodes. For the experimental setup described, L is approximately 5 µm, since within this distance we consider most of the contribution to the velocity (as shown in Fig. 2), and

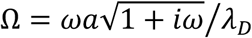

is the peak frequency at the scale of the charging time.

## RESULTS AND DISCUSSION

After these assumptions, from Eq. 1, the total capacitance can be described as the sum of capacitances linked in parallel and associated to the area composed of DNA strands (hybridized or not) and ions.

Referring to Fig. 4, the maximum percentage variation of C_d_ upon hybridization, C_d%_, for a given value of [cDNA], can be expressed as a function of the final (quantified after complete hybridization, 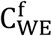) and initial (before hybridization, 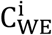) capacitance measured at the functionalized Working Electrode (WE):

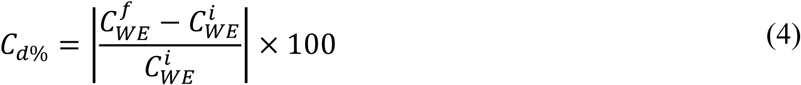

with

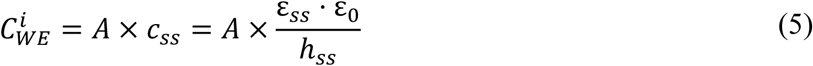

and

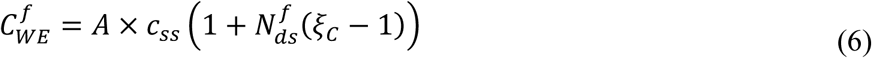

where *c*_*ss*_ (nF cm^−2^) is the density of capacitance measured at the ssDNA-SAM WE.

**FIGURE 4.**
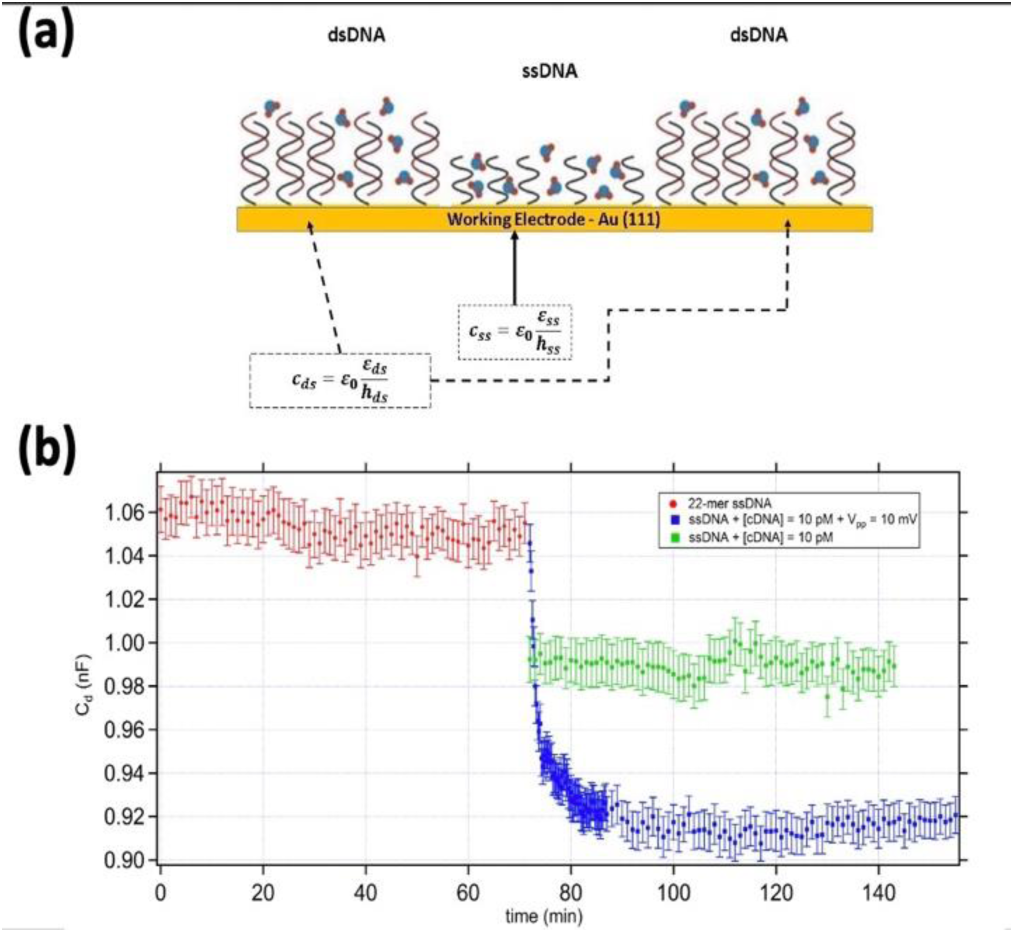
(A) Idealization of the electrode/electrolyte interface after a not complete hybridization process. (B) Hybridization of a Low Density (LD) ssDNA with an applied potential 10 mV in a buffer solution of 100 mM KCl performed by adding 10 pM complementary DNA.

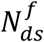 is the fraction of the total strands hybridized at the equilibrium. Obviously, it is function of [cDNA] and can be estimated by Sips model well described in (31, 32).

If h and ε are the thickness and the dielectric constant of the molecular layer before (ss) and after (ds) the hybridization process, we can fix the helpful fitting parameter:

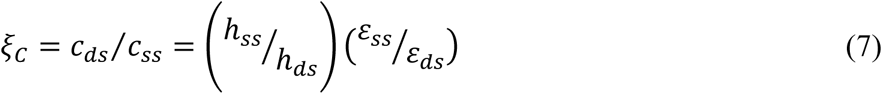

The amount of hybridized analyte^7^, on the gold surface, can be described by the differential equation resulting from the Langmuir adsorption model^15^ and that can be analytically solved in our experimental conditions:

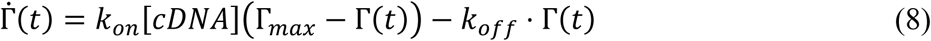

here, Γ(*t*) is the bound target coverage (molecules cm^−2^) for a given value of [cDNA], at a fixed temperature on the same probe film.

Rearranging the Eq. 4 in terms of Eqs. 2 and 3 and taking into account the Sips model (32), we found out the equation that models the charge density change at the interface:

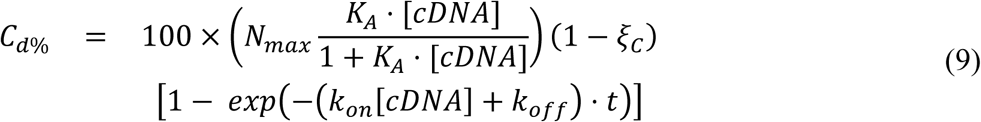

where 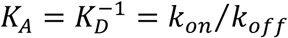 is the affinity constant for DNA hybridization and *k*_*on*_, *k*_*off*_ are the adsorption and desorption rate constants, respectively. N_max_ is the maximum number of hybridized sites for [cDNA].

By considering now the mass transport due to AC electroosmosis, we can assume an excess of targets ([T]_exc_, with respect to their initial value [T]_0_) in the formation rate of hybrids, [H], in the Langmuir adsorption model:

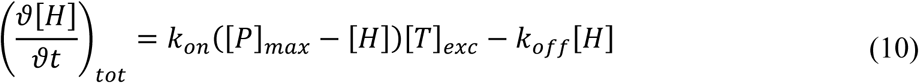

here, [*T*]_*exc*_ = [*T*]_0_ + [*T*]_*ACEO*_ and [*P*]_*max*_ is the maximum (initial) number of free probes on the surface.

Assuming no hybrids initially, the analytical solution of this differential equation is:

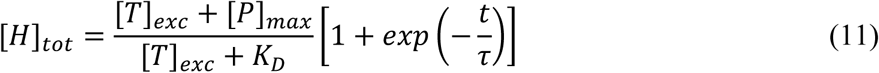

where 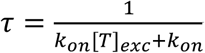.

In conclusion, by considering that at the equilibrium, the fraction of hybridized probes tends to 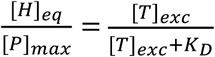, the percentage change of C_d_ can be estimated as:

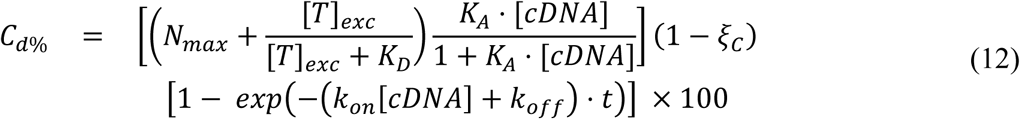

In both models (Eqs. 9 and 12), N_max_ is set equal to 0.49 and measure the maximum number of hybridized sites in a *suppressed hybridization regime* like the one defined by Wong et al. (10).

The Fig. 5 shows the simulation of our device with [1 pM to 100 nM] of cDNA strands in a 100 mM KCl-doped aqueous solution under an AC voltage of 10 mV. Here, we compare C_d%_ experimentally obtained and well fitted 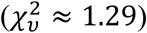 by a Hill Equation, (*K* ≅ 2.8 ± 0.7 nM) with the C_d%_ estimated via computational model including the excess of target due to the ACEO (*K*_*D*_ ≅ 3.46 nM, black fit) using the radius of gyration (≈ 1.78 nm, (29)) as radius of the DNA strand.

**FIGURE 5.**
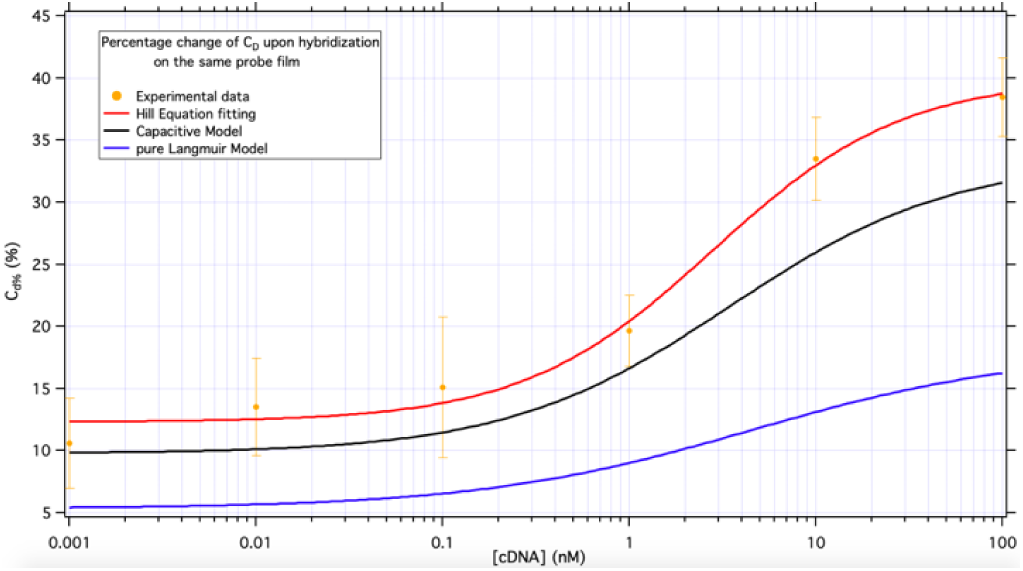
C_d%_ experimentally (orange markers) obtained as a function of [cDNA] dissolved in 100 mM KCl compared with the values got via two theoretical models given by the Eqs. 9 (blue curve) and 12 setting the radius of DNA strand equal to its radius of gyration (black fit). All the data are estimated in the same operating conditions. The red curve is the Hill equation that best fit our experimental data.

In Fig. 5, the blue curve represents the simulation performed in a pure Langmuir model (*K*_*D*_ ≅ 4.54 nM, Eq. 9), excluding therefore the excess of matter carried by ACEO. All the experiments and numerical simulations are performed in the same operating conditions.

In this study, the resulting velocity has been found to strongly depend on the input frequency, the electrical properties of the double layer and the applied electric field, as expected from the bibliographic survey.

The discrepancy between the theoretical prediction of AC electroosmosis and the experimental EIS results, suggest the need for more accurate handling of the electric double layer. The modelling of ACEO as a slip boundary condition based on the tangential component of the potential gradient across the electrode surface (Eq. 2) continue to be a valid approach, but the method we determined the potential drop across the electrical double layer may not be accurate for all conditions.

The computational solutions here show that, although polarizable particles can be moved using non-uniform electrical fields, the electrokinetiks forces are not the only forces acting on the strands. The total force on any particle is given by the sum of many forces including sedimentation, Brownian, dielectrophoretic and hydrodynamic forces; the latter arising from viscous drag on the particle.

It suggests that, whilst computational studies can model the phenomena occurring at the electrode/electrolyte interface and be used to optimize the performances of our biosensor, experimental data will be need to determine the practical conditions under which these phenomena occur.

## SUPPORTING MATERIAL

Supporting Material can be found online at

The role of the applied potential in reducing hybridization time, computation treatment of DNA molecule, Di-Electrophoresis (DEP) force calculation, AC-ElectroOsmosis (ACEO) driving and quantification, other useful values.

## AUTHOR CONTRIBUTIONS

P.C., S.D.Z. and Y.Y. conceived the experiments and interpreted data; P.C. performed the numerical simulations; P.C. and S.D.Z. performed the experiments and analyzed the data; P.C. and S.D.Z. fabricated and characterized the electrodes; P.C., S.D.Z., V.T. and Y.Y. wrote the paper. All the authors revised and approved the manuscript.

## ACKNOWLEDGMENTS

This work was supported by Associazione Italiana per la Ricerca sul Cancro (AIRC cinque per mille, no 12214) and FIRB 2011 grant RBAP11ETKA “Nano-technological approaches towards tumor theragnostic”.

We are grateful to Alessandro Bosco, L. Casalis, Pietro Parisse for a critical reading of the manuscript. P.C. thanks FNF @ IOM-CNR in Trieste for support in device fabrication.

